# Establishment of Tuberculosis Meningitis Mouse Model Using Aerosol Route of *W. Beijing* HN878 strain

**DOI:** 10.1101/2023.07.23.550249

**Authors:** Bhagwati Khatri, Sara Goulding, Vicky Rannow, Belinda Dagg, Mei Mei Ho

## Abstract

In this study, we report the development of a tuberculosis meningitis (TBM) mouse model using the HN878 strain via the aerosol route. Three genetically different strains of mice, CB6F1, C57BL/6 and BALB/c mice, were used to determine the dissemination of HN878 in the brain. The 8 x 10^8^ CFU/ml of HN878 dose was used to infect CB6F1 mice and deposited approximately 3.8 (0.07, SD) log_10_ CFU in the lungs. The burden of HN878 in the brain of the control group (administered saline) after approximately 16/17 days post-infection for CB6F1, C57BL/6 and BALB/c were 4.00 (0.47 SD), 3.79 (0.27, SD) and 2.12 (0.41, SD) log_10_ CFU/brain, respectively. The log_10_ CFU/brain in the BCG vaccinated CB6F1, C57BL/6 and BALB/c mice were 1.05 (0.61, SD), 2.13 (0.33, SD) and 1.42 (0.38, SD) respectively, which, if compared to the control groups, BCG vaccinated mice inhibited dissemination of HN878 in the brain by an impressive 2.94 (CB6F1), 1.66 (C57BL/6) and 0.69 (BALB/c) log_10_ CFU/brain reduction. In conclusion, we have established a relatively inexpensive TBM mouse model using an aerosol, a natural route of infection, which will advance research in understanding TBM dissemination to the brain, and preclinical tuberculosis vaccine/drug discovery/drug regimens against TBM.

## 1 Introduction

In 2021, approximately 10.6 million people fell ill due to tuberculosis (TB) and 1.6 million people died of TB (WHO, 27 Oct 2022). From the retrospective region-specific studies, of all people with TB, 0.3 – 4.9% have the deadliest form of TB, tuberculosis meningitis (TBM) (Seddon et al., 2019, Huynh et al., 2022). The actual incidence rate of TBM is not known and is likely to be substantially under-reported due to the difficulties in diagnosis and inadequate surveillance. However, in settings with a high tuberculosis burden, TBM, disproportionately affects young children and individuals with HIV (Seddon et al., 2019). Bacillus Calmette-Guérin (BCG) vaccine confers the most protection against TBM, especially in neonates and school-age children, with an efficacy of 70%. Nevertheless, the vaccine efficacy is much lower in adults (Mangtani et al., 2014), accounting for most TB transmission. In 2019 (du Preez et al., 2019), the impact of the global shortages of the BCG vaccine linked to the rise of TBM in children was reported. Furthermore, with the combined complications of HIV-1 co-infection and multi-drug resistance in TBM, the mortality is close to 100% (Marais et al., 2011).

This debilitating TBM disease is an extrapulmonary manifestation of TB caused by *Mycobacterium tuberculosis* (Mtb). TBM affects the brain and spinal cord, leading to complications such as hydrocephalus (buildup of fluid in the brain), vasculitis (inflammation of the blood vessels), encephalitis (inflammation of the brain) and inflammation of the meninges (Chin, 2014). Much of the research has advanced in understanding the pathogenesis, diagnostics and drug treatment, but TBM research in humans is complex. Multiple factors are involved in the difficulty of conducting TBM research in humans (Huynh et al., 2022, Davis et al., 2019, Davis et al., 2020, Wilkinson et al., 2017), such as the inability to accurately determine the precise time a patient was infected; the mechanism of brain injury and their relation to inflammation; the moment and conditions under which bacteria spread to the central nervous system (CNS); the impossibility of obtaining CNS samples from patients, which is vital to understand immune markers in the onset of TBM. Furthermore, the next-generation TB vaccine or drug selection and testing are performed in the preclinical animal models against pulmonary TB and lacks data on the prevention of dissemination of TB in the brain. Establishing the relevant TB animal model would aid further advancement in understanding pathogenesis, vaccines or drug selection to advance in clinical trials.

Animal models are indispensable in progressing TBM research to address the pertinent research gaps mentioned above. Although TBM animal models exist, the Mtb infection route in animals mainly utilises invasive procedures in establishing infection in the brain. For example, the first models of TBM were established in the early 20^th^ century using dogs (Manwaring, 1912), rabbits (Kasahara, 1924) and guinea pigs (Shope and Lewis, 1929). TBM was established in the brain of these early animal models via intracerebral Mtb inoculation. Subsequently, over the years, various routes of Mtb infection in guinea pigs were performed, such as subcutaneous/intracerebral (Burn and Finley, 1932), intravenous/intrathecal (Rich, 1933) and an aerosol (Skerry et al., 2013). The rabbit model of TBM has been developed using Mtb infection routes of intrathecal (Tsenova et al., 2006) and intracisternal (Tsenova et al., 2005). The TBM mouse model used different genetic backgrounds of mice, and Mtb infection was performed by various routes, such as intracerebral (Zucchi et al., 2013, Lee et al., 2009), intracranial (Lee et al., 2009), intravenous (Be et al., 2008) and intratracheal (Hernandez Pando et al., 2010). The guinea pig is the only model with the natural route of infection (Skerry et al., 2013); however, this model has limitations, such as the housing capacity in a containment level III environment, the cost of animal experiments, limited facilities to infect guinea pigs via aerosol route and limited reagents. Therefore, we sought to establish a relatively inexpensive, easy-to-adopt mouse model of TBM via the natural route of aerosol infection.

In the study, we report the development of the TBM mouse model in three different genetic background strains of mice, CB6F1, C57BL/6 and BALB/c. The whole body exposure (aerosol route) using *W. Beijing* HN878 (HN878) at a dose of 8 x 10^8^ colony forming unit (CFU)/ml was selected and used for infecting mice. The study tested the efficacy of the BCG vaccine against HN878 infection and observed a significant reduction in the dissemination of HN878 to the brain for all strains of mice compared to the control mice. Furthermore, we observed clinical symptoms/body weight loss in the non-vaccinated CB6F1, C57BL/6, but not BALB/c mice. To our knowledge, this will be the first report using the aerosol route of HN878 infection in establishing a TBM mouse model using CB6F1, C57BL/6 and BALB/c strains of mice.

## 2 Materials and Methods

### 2.1 Ethics, animals and study groups

All animal procedures were performed in accordance with the UK Home Office (Scientific Procedures) Act 1986; under appropriate licences. The Animal Welfare and Ethics Review Body at the South Mimms site of MHRA approved the study protocol.

CB6F1, C57BL/6 and BALB/c mice, female, 6-8 weeks or 10-12 weeks old, with approximately >15 gm and >20 gm body weight, respectively, were obtained from the Specific Pathogen-Free facilities at the Charles River UK. Ltd. All animals were housed in appropriate containment level II (BCG vaccinated) and containment level III (Mtb infected) facilities at the South Mimms site, MHRA. Throughout the experiment, the animal technicians monitored the health and well-being of all mice daily as per the Animals in Science Regulation Unit (Home Office, 26 Oct 2022), an act for the welfare of animals. All mice were weighed before HN878 infection and subsequently weighed daily post-infection (PI), except in one experiment using CB6F1 mice (section 3.1), where mice were observed instead of weighed. The criteria for terminating the mice during the experiments were if the mice nearing 20% body weight loss and/or onset of clinical symptoms, such as laboured/rapid breathing, piloerect coat, hunched posture and lethargy. The mice were terminated by the schedule 1 method, followed by confirmation of death. The frequency of animal checking increased after indicating body weight loss (more than 15% body weight). The study was performed in compliance with the ARRIVE guidelines. A high protein diet (DietGel® Boost from Clear H_2_O) was administered to CB6F1 mice in one of the experiments (section 3.3). The study groups were five mice per group for all experiments. The study groups for the CB6F1 mice (section 3.2) were; a saline-administered group (control group) and two BCG vaccinated groups (BCG groups), BCG and BCG-group 2. The C57BL/6 and BALB/c mice (section 3.2) had one control group and one BCG group. The age of all mice at the time of HN878 infection was 10-12 weeks.

### 2.2 BCG vaccination and HN878 challenge

CB6F1, C57BL/6 and BALB/c mice were intradermally vaccinated with the WHO Reference Reagent of BCG vaccine, Danish 1331 sub-strain preparation (NIBSC code 07/270; https://www.nibsc.org/products/brm_product_catalogue/detail_page.aspx?catid=07/270). The lyophilised BCG Danish 1331 reference reagent was diluted in sterile saline with the aim of administrating a BCG dose of 3-4 x 10^4^ CFU per mouse. The BCG dose was checked by plating in duplicate onto 7H11 agar plates (DIFCO) supplemented with Oleic Acid Dextrose Catalase (OADC, BD), and CFU/ml was estimated at four weeks post-incubation at 37^0^C. All vaccinated mice received an expected dose of BCG at 3-4 x 10^4^ CFU/mouse (data not shown).

For aerosol infection, we used the procedure described in our previous publication (Khatri et al., 2021). Briefly, HN878 infection inoculum was prepared from the HN878 frozen stock (PBS containing 50% glycerol), which was diluted in sterile distilled water to approximately 8 x 10^8^ CFU/ml and 1 x 10^8^ CFU/ml (for CB6F1 mice only, section 3.1). The HN878 infection inoculum was transferred in a Glass nebuliser-venturi (Glas‐Col) (Ordway and Orme, 2011, Chan et al., 2020) and mice were infected for 30 minutes using the whole body exposure chamber (Walkers, UK), which operates on a similar concept as the Madison infection chamber (Ordway and Orme, 2011). The CB6F1 mice (5/group) infected with the HN878 doses at 8 x 10^8^ CFU/ml and 1 x 10^8^ CFU/ml were used to estimate the log_10_ CFU/lung deposition, one-day post-infection (PI). The lungs of the mice were harvested, homogenised, and serially diluted homogenates were plated in duplicate onto 7H11 agar (DIFCO) supplemented with OADC (BD). Bacterial colonies were enumerated 3-4 weeks post-incubation at 37^0^C and presented throughout as log_10_ CFU/organ.

For all experiments, unless specified, mice were stored at -20^0^C after termination and thawed later for organ harvesting. After the mice were thawed, the lungs, brain and spleen were harvested, homogenised and log_10_ CFU/organ was enumerated as described above.

### Statistical analyses

All data were analysed using the GraphPad Prism 8 software (Graph Pad, USA) using a student’s t-test or one-way ANOVA with Dunnett’s post-test statistical analyses. The HN878 burden in the organs is presented as an average of five mice log_10_ CFU/organ ± standard deviation (SD). The following formulas were used for the log reduction and percentage reduction in the BCG group (log_10_ CFU/organ) compared to the control group:

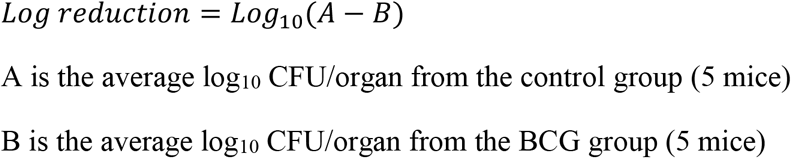

For converting Log reduction to the percentage reduction:

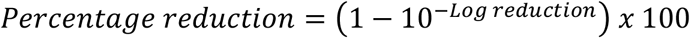

The percentage body weight change was calculated using the following formula:

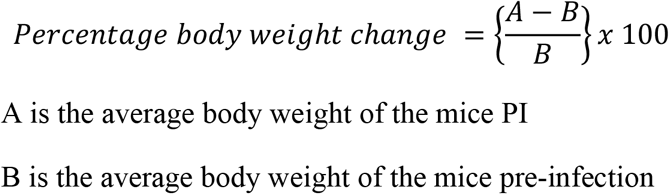

A ρ value <0.05 was considered as a statistical difference and denoted in the figures with * ρ <0.05, ** ρ <0.005, *** ρ <0.001 and **** ρ <0.0001.

## 3 Results

### 3.1 Selection of HN878 infection dose

For the standard aerosol Mtb infection model (deposits ∼100 CFU/lungs), the lungs and spleen, but not the brain, are generally harvested after four weeks of infection to determine Mtb burden. The Mtb dissemination to the brain via a standard aerosol Mtb infection is not well defined. Therefore, to understand the Mtb burden in the brain of a standard aerosol Mtb infection dose of (6-7 x 10^6^ CFU/ml), the brains of Mtb-infected CB6F1 and C57BL/6 mice were harvested from some of the provisional mice experiments from our lab. All mice were aerosol-infected with the standard Mtb dose of 6 – 7 x 10^6^ CFU/ml, which approximately deposits 100 CFU/lungs per mouse after one-day PI. Various Mtb strains from lineages 2, 3 and 4 were used for infection. The Mtb lineage 2 strains were; HN878; GC1237; a couple of clinical isolates from Kenyan individuals, Beijing 1521 and clade T2. From the lineage 3 – CAS1-Kili strain, lineage 4 – clade T1 (a clinical isolate from a Kenyan individual) and a reference strain H37Rv were used to infect mice. The fresh brains were harvested at four weeks or 12 weeks PI from all mice infected with various Mtb strains. The CB6F1 (Fig. 1A) and C57BL/6 mice (Fig. 1B), at four weeks PI, displayed variability within the groups for Mtb burden in the brain, with most of the mice showing less than 2 log_10_ CFU/brain. For the C57BL/6 mice (Fig. 1B), at 12 weeks PI, Mtb burden in the brain of mice infected with HN878 and GC1237 displayed an average of 1.7 ± 0.47 and 1.4 ± 0.28 log_10_ CFU/brain, respectively. The rest of the mice harvested at 12 weeks PI for various Mtb strains either showed no growth of Mtb or less than 2 log_10_ CFU/brain.

**Figure 1:**
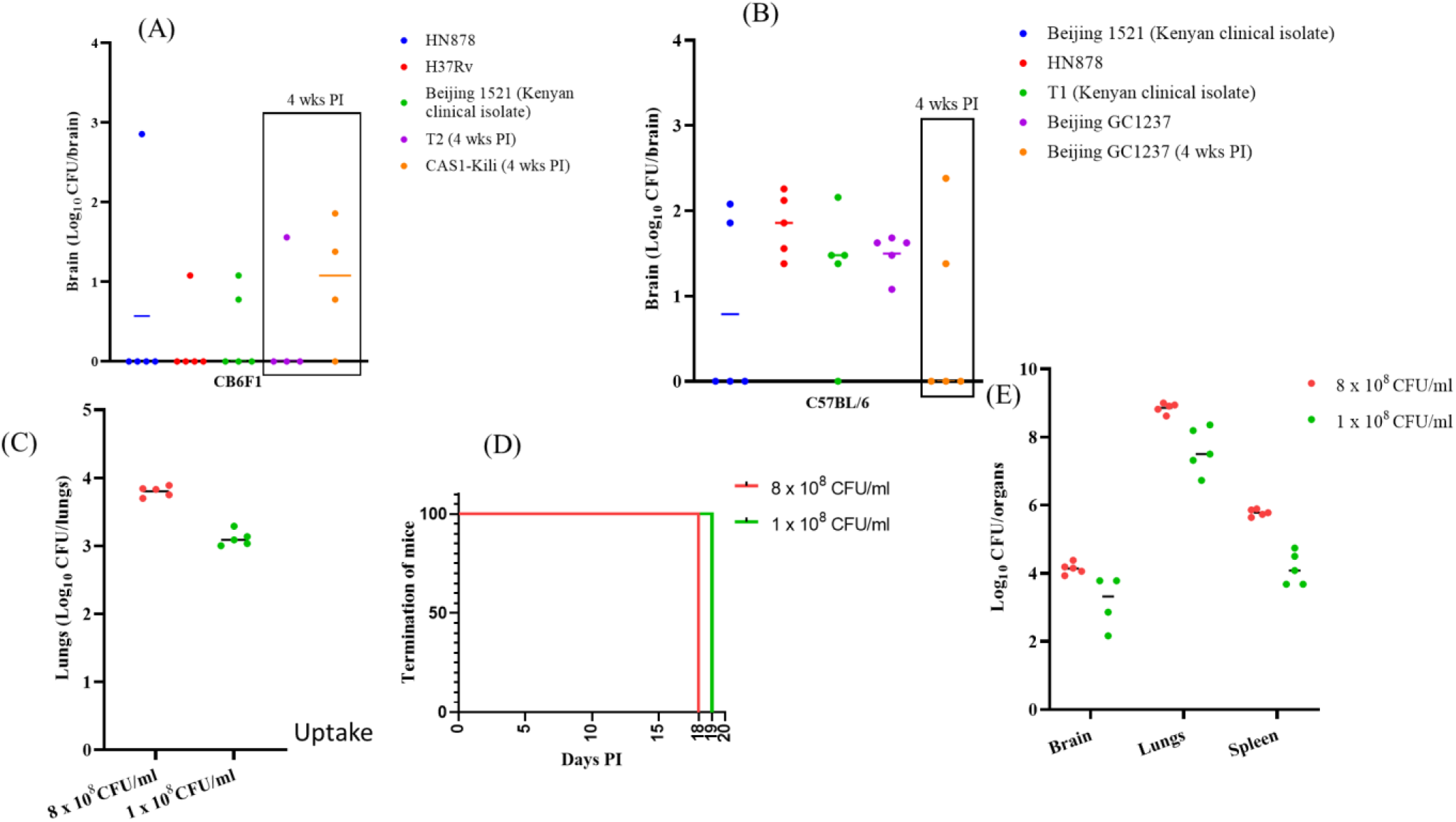
Evaluating standard infection dose using various Mtb strains and selection of HN878 challenge dose. **(A)**, CB6F1 mice infected with HN878, H37Rv, clinical isolates from Kenyan individuals (1521, T2 clade and CAS1-kili) with a standard challenge dose of 6-7 x 10^6^ CFU/ml (∼100 CFU/mouse lung deposition after one-day of infection). The fresh brains were harvested from the infected CB6F1 mice for log_10_ CFU/brain determination after 12 weeks (HN878, H37Rv & 1521) and four weeks (T2 clade and CAS1-Kili) PI. **(B)**, C57BL/6 mice infected with HN878, GC1234, clinical isolates from Kenyan individuals (1521 and T1 clade) with a standard infection dose as of (A). The fresh brains were harvested from the infected C57BL/6 mice for log_10_ CFU/brain determination after 12 weeks (HN878, GC1234, 1521 and T1 clade) and four weeks (GC1234) PI. **(C)**, CB6F1 mice were infected with two doses of HN878, 8 x 10^8^ CFU/ml and 1 x 10^8^ CFU/ml, via the aerosol route. After one day PI, the brains of the CB6F1 mice (5/group) from HN878 doses, 8 x 10^8^ CFU/ml and 1 x 10^8^ CFU/ml, were harvested for log_10_ CFU/brain determination. **(D)**, the termination of the CB6F1 mice infected with both HN878 doses displayed clinical symptoms by 18 days (8 x 10^8^ CFU/ml) and 19 days (1 x 10^8^ CFU/ml). **(E)**, the brain, lungs and spleen from the CB6F1 mice were harvested 18 days PI from the HN878 dose of 8 x 10^8^ CFU/ml and 19 days PI of 1 x 10^8^ CFU/ml for log_10_ CFU/brain determination. The mean for all graphs, except graph (D), is represented with a horizontal line.

Therefore, to aim to increase the burden of Mtb in the brain, two doses of HN878 infection inocula, 8 x 10^8^ CFU/ml and 1 x 10^8^ CFU/ml, were used to infect CB6F1 mice. After one day, the lungs from one group (uptake mice) of CB6F1 mice (5/group) were harvested to determine the amount of HN878 deposited in the lungs. The average of 3.8 ± 0.07 and 3.1 ± 0.11 log_10_ CFU/lungs (Fig. 1C) was deposited in the lungs of uptake mice for HN878 doses at 8 x 10^8^ CFU/ml and 1 x 10^8^ CFU/ml, respectively. The other groups of CB6F1 mice, which were infected with HN878 at 8 x 10^8^ CFU/ml and 1 x 10^8^ CFU/ml, started clinical symptoms of laboured/rapid breathing, piloerect coat, hunched posture, lethargy and were terminated by schedule 1 method at days 18 and 19 PI, respectively (Fig. 1D). The brain, lungs and spleen were harvested to determine HN878 burden per organ. For mice infected with the HN878 dose at 8 x 10^8^ CFU/ml, Mtb burden in the organs at 18 days PI was 4.14 ± 0.17 log_10_ CFU/brain, 8.86 ± 0.15 log_10_ CFU/lungs and 5.78 ± 0.1 log_10_ CFU/spleen (Fig. 1E). For mice infected with the HN878 dose at 1 x 10^8^ CFU/ml, Mtb burden in the organs at 19 days PI was 3.14 ± 0.79 log_10_ CFU/brain, 7.62 ± 0.67 log_10_ CFU/lungs and 4.14 ± 0.48 log_10_ CFU/spleen (Fig. 1E). The brain data of one mouse infected with HN878 dose of 1 x 10^8^ CFU/ml was excluded from Fig. 1E due to the contamination issue in the 7H11 plates.

The HN878 dose of 8 x 10^8^ CFU/ml resulted in a six times higher HN878 burden in the brain than the 1 x 10^8^ CFU/ml. In addition, both HN878 doses induced clinical symptoms in the CB6F1 mice, followed by schedule 1 termination almost similar days PI (18 days and 19 days). Therefore, the HN878 dose at 8 x 10^8^ CFU/ml was chosen for all the subsequent experiments.

### 3.2 BCG vaccination significantly reduces the HN878 dissemination in the brain for CB6F1, C57BL/6 and BALB/c mice

Next, to examine the BCG vaccine effect on the HN878 dissemination in the brain and the BCG-induced protection in different mice genetic backgrounds, three strains of mice were used CB6F1, C57BL/6 and BALB/c mice. All mice were vaccinated with BCG at 3-4 x 10^4^ CFU/mouse (BCG group) or administered saline (control group). After four weeks of BCG vaccination or saline administration, mice were infected with the HN878 dose at 8 x 10^8^ CFU/ml via the aerosol route. Fig. 2A-C shows the percentage body weight changes for CB6F1, C57BL/6 and BALB/c. The control group of CB6F1 and C57BL/6 mice (Fig. 2A&B) started weight loss from 14 and 13 days PI, respectively. The control group of CB6F1 mice (Fig. 2A), at 16 days PI, was nearing 20% body weight loss, and the mice were terminated on welfare grounds. Similarly, in the control group of C57BL/6 mice (Fig. 2B), the body weight loss at 17 days PI was nearing 20% followed by mice termination. The BCG groups of CB6F1 (Fig. 2A) and C57BL/6 (Fig. 2B) mice were culled simultaneously when the counter-control groups were terminated to compare the HN878 burden in the organs. The BCG groups displayed no clinical symptoms; in addition, for the CB6F1 mice (Fig. 2A), another set of BCG group (BCG – group 2) was culled at 24 days PI to evaluate the dissemination of HN878 in the brain. Interestingly, the control group of BALB/c mice (Fig. 2C) displayed no weight loss until 16 days PI, but at 17 days PI, a drop in body weight was observed. Although BALB/c mice did not show any clinical symptoms, they were terminated to determine the HN878 burden in the organs at approximately the same time point as CB6F1 and C57BL/6 mice.

**Figure 2:**
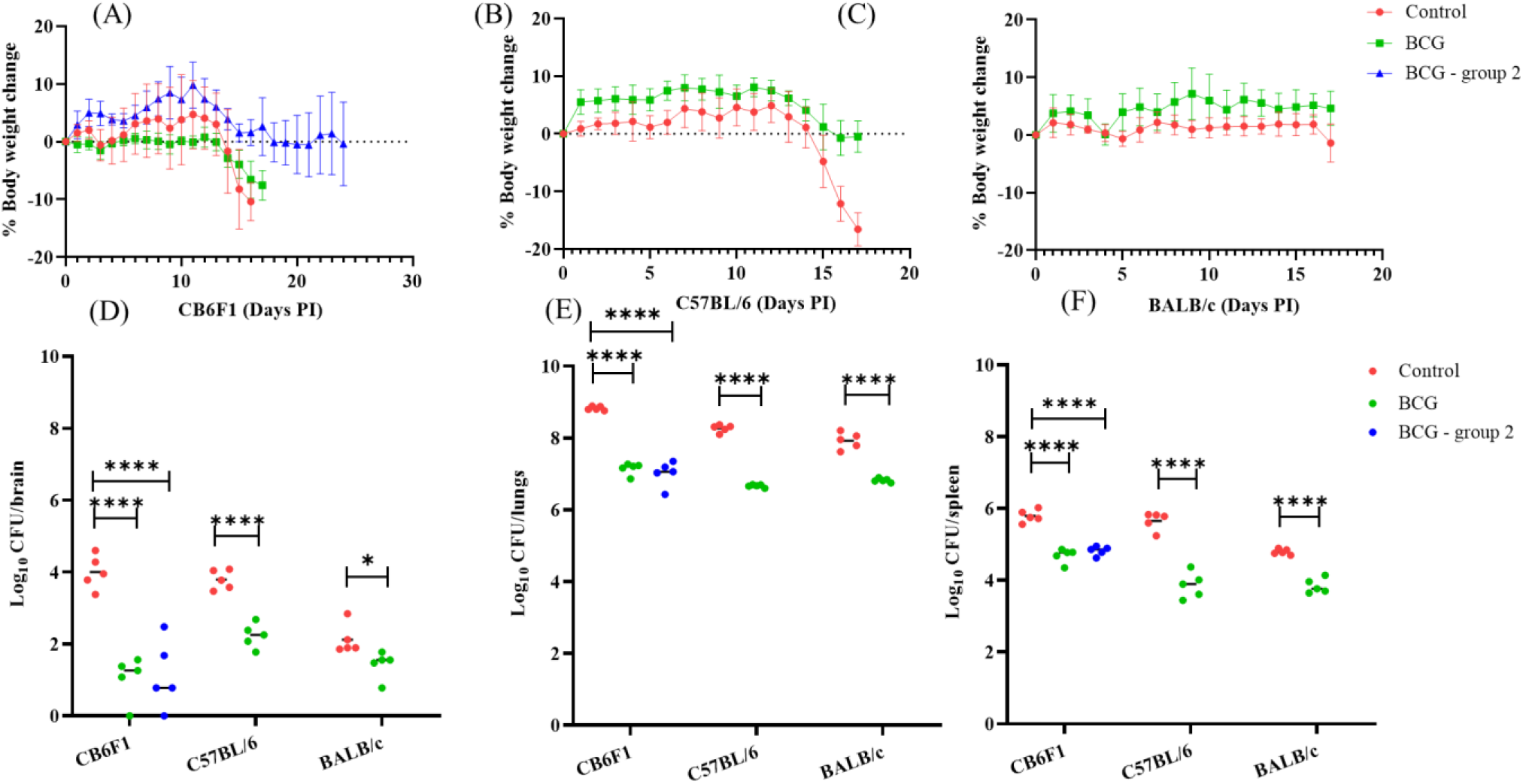
Body weight changes after HN878 infection and BCG vaccination in CB6F1, C57BL/6 and BALB/c mice. **(A)**, the percentage body weight change for CB6F1 mice PI at 8 x 10^8^ CFU/ml. The BCG group was culled one day after the control group (16 days) and the BCG-group 2 was culled at 24 days PI. **(B)**, the percentage body weight change for C57BL/6 mice PI at 8 x 10^8^ CFU/ml. **(C)**, the percentage body weight change for BALB/c mice PI at 8 x 10^8^ CFU/ml. (D), the HN878 burden in the brain (log_10_ CFU/brain) for CB6F1, C57BL/6 and BALB/c mice. All strains of mice had control and BCG groups, except CB6F1, which had an additional BCG-group 2. (E), the HN878 burden in the lungs (log_10_ CFU/lungs) of CB6F1, C57BL/6 and BALB/c mice. (F), the HN878 burden in the spleen (log_10_ CFU/spleen) of CB6F1, C57BL/6 and BALB/c mice. Statistical analysis for **(A), (B)** and **(C)**, the data are the mean percentage body weight change ± SD for CB6F1, C57BL/6 and BALB/c mice, respectively. **(D), (E)** and **(F)** dots represent each mouse’s log_10_ CFU/organ, with the mean as a horizontal line. The Dunnett’s multiple comparison test, one-way ANOVA was used for CB6F1 mice (BCG and BCG-group 2) and student’s t-test for C57BL/6 and BALB/c mice to determine the statistical significance between the vaccinated groups compared to the control group. * ρ <0.05 and **** ρ <0.0001.

The brain, lungs and spleen from all groups of CB6F1, C57BL/6 and BALB/c mice (Fig. 2D-F) were harvested for log_10_ CFU/organ determination. The HN878 burden in the brains of the control groups for CB6F1, C57BL/6 and BALB/c mice (Fig. 2D) were log_10_ CFU/brain, 4.00 ± 0.47, 3.79 ± 0.27 and 2.12 ± 0.41, respectively. For the BCG groups, the log_10_ CFU/brain for CB6F1, C57BL/6 and BALB/c mice were 1.05 ± 0.61 & 1.14 ± 0.95 (BCG & BCG – group 2), 2.13 ± 0.33 and 1.42 ± 0.38, respectively. The significant log_10_ reduction in the BCG groups compared to the control groups for CB6F1, C57BL/6 and BALB/c mice were 2.94 & 2.86 (BCG & BCG – group 2), 1.66 and 0.69, respectively, which equates to percentage reduction of 99.88% & 99.86% (for BCG & BCG-group 2), 97.21% and 79.66%, respectively.

The HN878 burden in the lungs for the control groups of CB6F1, C57BL/6 and BALB/c mice (Fig. 2E), the log_10_ CFU/lungs were – 8.83 ± 0.05, 8.27 ± 0.10 and 7.92 ± 0.23, respectively. For the BCG groups of CB6F1, C57BL/6 and BALB/c mice (Fig. 2E), the log_10_ CFU/lungs were 7.14 ± 0.16 & 7.08 ± 0.20 (BCG & BCG – group 2), 6.66 ± 0.04 and 6.82 ± 0.05, respectively. The significant log reduction in the BCG groups compared to the control groups for CB6F1, C57BL/6 and BALB/c mice were 1.68 & 1.75 (BCG & BCG – group 2), 1.60 and 1.10, respectively, which equates to percentage reduction 97.92% & 98.20% (for BCG & BCG-group 2), 97.50% and 92.13%, respectively.

Furthermore, the HN878 burden in the spleen for the control groups of CB6F1, C57BL/6 and BALB/c mice (Fig. 2F), the log_10_ CFU/spleen were – 5.79 ± 0.18, 5.65 ± 0.25 and 4.79 ± 0.08, respectively. For the BCG groups of CB6F1, C57BL/6 and BALB/c mice (Fig. 2F), the log CFU/spleen were 4.69 ± 0.20 & 4.82 ± 0.13 (BCG - group 1 & BCG – group 2), 3.87 ± 0.36 and 3.84 ± 0.20, respectively. The significant log_10_ reduction in the BCG groups compared to the control groups for CB6F1, C57BL/6 and BALB/c mice were 1.10 & 0.97 (BCG & BCG – group 2), 1.79 and 0.94, respectively, which equates to percentage reduction of 92.06% & 89.20% (for BCG & BCG-group 2), 98.37% and 88.75%, respectively.

### 3.3 The HN878 burden in the brain of 14 days infected CB6F1 mice with a high protein diet administration

The welfare of the animals is critical to minimise suffering during the experiments. Therefore, an experiment was set up to examine a high protein diet effect on the dissemination of HN878 to the brain of CB6F1 mice 14 days PI. The CB6F1 mice were BCG vaccinated (BCG group), or saline was administered (control group). After four weeks of BCG vaccination, all mice were challenged with the HN878 at 8 x 10^8^ CFU/ml. At seven days PI, some mice from the control group displayed a slight decline in body weight (Fig. 3A). Therefore, all mice from both groups were supplemented with a high-protein diet from seven days PI and were terminated at 14 days PI with no clinical symptoms.

**Figure 3:**
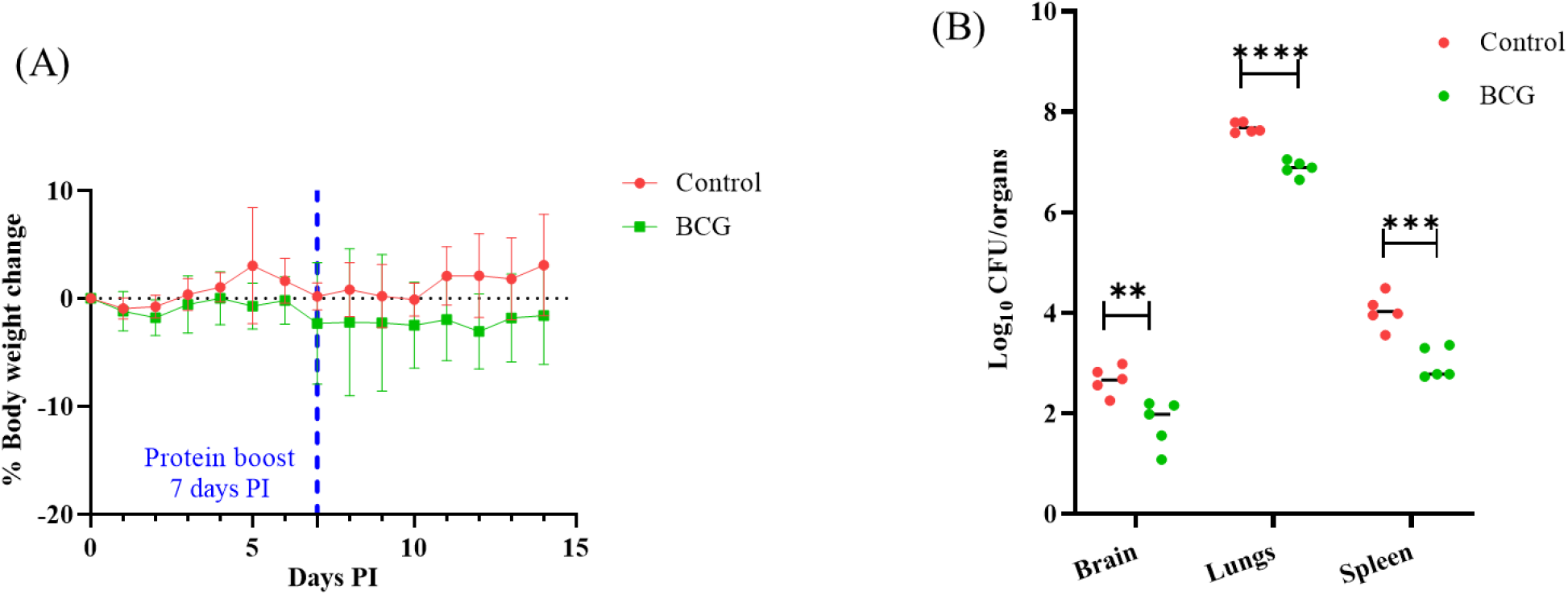
High protein diet administration and 14 days PI harvest for CB6F1 mice. **(A)**, CB6F1 mice infected with HN878 at 8 x 10^8^ CFU/ml. The percentage body weight change for CB6F1 mice (5/group) was plotted as mean ± SD for control and BCG groups. The blue vertical line shows the administration of a protein diet initiated on the seventh day PI until the termination of both groups on 14 days PI. **(B)**, dots represent each mouse’s log_10_ CFU/organ, with the mean as a horizontal line. Student’s t-test was performed to determine the statistical significance between each organ of vaccinated groups compared to the control group. ** ρ <0.005, *** ρ <0.001 and **** ρ <0.0001.

The brain, lungs and spleen were harvested to determine HN878 CFU/organ (Fig. 3B). The average log_10_ CFU/organ for the control group was 2.64 ± 0.29 (brain), 7.68 ± 0.10 (lungs) and 4.02 ± 0.38 (spleen). For the BCG group, the average log_10_ CFU/organ was 1.79 ± 0.47 (brain), 6.88 ± 0.01 (lungs) and 2.99 ± 0.31 (spleen). The log reduction in the organs of BCG groups compared to the control groups for CB6F1 was 0.86 (brain), 0.80 (lungs) and 0.87 (spleen). The conversion of log reduction to the percentage reduction of HN878 burden in the BCG group, when compared to the control group, would equate to 86.26% (brain), 84.15% (lungs) and 86.36% (spleen).

## 4 Discussion

Although TBM research has advanced substantially, some research is impossible in humans due to ethical issues in collecting samples. Therefore, a much-needed TBM mouse model using the natural infection route will aid in advancing TBM research. Many important TBM questions exist (Huynh et al., 2022) - what are the underlying mechanisms of cerebral infarctions, tuberculomas, and hydrocephalus? What is the mechanism of brain injury and its relationship with neuroinflammation? How to improve the sensitivity of pathogen-based diagnostic testing? What is the optimal dose of rifampicin in the treatment of TBM? Can the duration of TBM therapy be shortened? What are the optimal dose and duration of adjuvant therapy?

Establishing the TBM mouse model will be an invaluable tool to answer some of the important TBM questions mentioned above. Furthermore, the TBM mouse model has the potential to serve as an indispensable tool in testing anti-TBM drugs and next-generation TB vaccines. BCG vaccine confers the most protection against TBM. The next-generation TB vaccine testing is generally performed in preclinical animal models, and the vaccines’ effectiveness is mainly assessed in the lungs. With the establishment of our TBM mouse model, the next-generation TB vaccine efficacy can be assessed simultaneously for pulmonary TB and TBM, thereby allowing the selection of TB vaccines against both, TBM and pulmonary TB.

The selection of the infecting HN878 doses at 8 x 10^8^ CFU/ml or 1 x 10^8^ CFU/ml was performed in the CB6F1 mice and was based on determining the HN878 burden in the brain. The HN878 dose at 8 x 10^8^ CFU/ml had one log_10_ higher HN878 burden in the brain (4.14 ± 0.17 log_10_ CFU/brain) compared to the 1 x 10^8^ CFU/ml (3.14 ± 0.79 log_10_ CFU/brain). In addition, both HN878 doses, 8 x 10^8^ CFU/ml and 1 x 10^8^ CFU/ml, displayed clinical symptoms in mice and had to be culled on welfare grounds around similar days of 18 and 19 days, respectively. Furthermore, mice in the 1 x 10^8^ CFU/ml dose displayed overall variable results in the organs when compared to the 8 x 10^8^ CFU/ml dose. Based on all the data observations for HN878 dose selection, the highest dose of the HN878 at 8 x 10^8^ CFU/ml was chosen for all the subsequent experiments.

Dissemination of HN878 in the brain was an encouraging step forward in establishing the TBM mouse model, but a few questions were pending in further testing the TBM model: Do the host genetic factors influence the degree of HN878 dissemination to the brain? Does the BCG vaccine prevent the HN878 dissemination to the brain? If so, what degree of protection does the BCG vaccine elicit? To answer these questions, we chose three different genetic backgrounds of mice, CB6F1, C57BL/6 and BALB/c mice. All strains of mice had at least two groups, the BCG vaccinated group and the control group, infected with HN878 at 8 x 10^8^ CFU/ml dose after four weeks of BCG vaccination. The CB6F1 and C57BL/6 mice, but not BALB/c mice, displayed nearing 20% body weight loss on 16 and 17 days PI, respectively and had to be culled on welfare grounds. However, although the BCG groups of CB6F1 and C57BL/6 mice displayed a drop in body weight after PI, they seemed to stabilise; for example, the CB6F1 mice, BCG group 2, which was culled at 24 days PI, displayed body weight loss from 11 days PI, but stabilised the weight after 16 days with some fluctuations until 24 days PI. Similarly, the BCG group of C57BL/6 mice displayed body weight loss by 11 days PI, but stabilised from 16 days PI and culled at 17 days PI at the same time when the control group of C57BL/6 mice was terminated.

The HN878 burden in the brains of CB6F1 (16 days PI) and C57BL/6 (17 days PI) mice from the control groups was 4.00 ± 0.47 and 3.79 ± 0.27 log_10_ CFU/brain, respectively. The BCG vaccine in CB6F1 mice prevented HN878 dissemination to the brain by around an impressive 99.88% (nearly three log reduction). The BALB/c mice had shown the least HN878 burden in the brain of control mice, 2.12 log_10_ CFU/brain and the BCG vaccine protection was around 79.66% (0.69 log reduction). In one of our published studies (Khatri et al., 2021), we demonstrated that the cytokines IL2, GM-CSF and CXCL13 in the supernatants of stimulated splenocytes were significantly inversely correlated in the lungs of HN878-infected CB6F1 mice, but not in BALB/c mice. The role of cytokines IL2, GM-CSF and CXCL13 in CB6F1 mice and their relation in disseminating Mtb to the brain would be an exciting area for further investigation.

The host genetic factors may be involved in the dissemination of Mtb in the brain. However, it cannot be ruled out that there might be a positive correlation between the amount of HN878 burden in the lung and its dissemination to the brain. For example, in Fig. 2D&E, the control group of CB6F1 mice had the most HN878 burden in the lungs (8.83 ± 0.05 log_10_ CFU/lungs) and in the brain (4.00 ± 0.47 log_10_ CFU/brain), followed by C57BL/6 (8.27 ± 0.10 log_10_ CFU/lungs and 3.79 ± 0.27 log_10_ CFU/brain) and BALB/c (7.92 ± 0.23 log_10_ CFU/lungs and 2.12 log_10_ CFU/brain).

Both the HN878 doses at 8 x 10^8^ CFU/ml or 1 x 10^8^ CFU/ml initiated CB6F1 mice body weight loss around similar time points and the mice had to be sacrificed on welfare grounds after developing visible distressing signs (Fig. 1D). The HN878 deposition in the brain and lungs of CB6F1 mice after 18 days PI for dose 8 x 10^8^ CFU/ml was approximately three and 17 times more compared to the 19 days PI of 1 x 10^8^ CFU/ml dose, respectively. The clinical symptoms in the CB6F1 mice displayed more physical distress signs than neurological symptoms in mice, such as ataxia or akinesia. This may suggest that the weight loss was the onset of a possible high TB burden in the lungs instead of TB in the brain.

The development of the TBM mouse model in three different genetic background strains of mice had its challenges, especially the inception of adverse effects, body weight loss in the mice, and the selection of the type of Mtb strain and dosage for infecting the mice. The selection of C57BL/6 and BALB/c mice has mainly been based on their availability and use in TB research. The CB6F1 mice are the hybrid of C57BL/6 and BALB/c mice used in our lab (Khatri et al., 2021) for the standard aerosol HN878 dose (∼100 CFU/mouse) studies. An initial assessment of the adverse effects was performed in the CB6F1 when challenged with two HN878 doses, 8 x 10^8^ CFU/ml and 1 x 10^8^ CFU/ml and for both doses, mice had to be culled due to clinical symptoms. However, the introduction of the protein diet to CB6F1 mice at seven days PI and harvesting organs at 14 days PI, unfortunately, failed to show a high HN878 burden in the brain (2.6 ± 0.28 log_10_ CFU/brain). This might be due to reasons; A, as discussed earlier, the positive correlation of HN878 burden in the lungs (7.68 ± 0.10 log_10_ CFU/lungs) and the possibility of simultaneous dissemination of HN878 in the brain (2.64 ± 0.29 log_10_ CFU/brain), Fig. 3B. B, possible that the HN878 is still multiplying in the brain at 14 days, which may reach 18 days PI peak of 4.14 ± 0.17 log_10_ CFU/brain. However, the effect of a protein diet in stabilising the control mice’s body weight could not be established due to the early termination of the experiment at 14 days instead of 17 days.

The development of the TBM mouse model is a significant step forward and will hasten in answering some of the pertinent questions, such as how Mtb disseminates from the lungs to CSF. What immune responses are involved in the onset of TBM? How to improve the diagnosis of TBM? Finally, developing a relatively inexpensive and reasonably easy-to-adopt TBM aerosol mouse model will be a crucial step forward in assessing the next-generation TB vaccines (assess the brain protection in parallel to the lungs), anti-TB drug testing and new drug regimens or combination therapy.

## 6 Author Contributions

Conceptualisation, Project Lead, Formal analysis, Data Interpretation, Visualisation, Writing-original draft; BK. Conducting experiments; SG, VR, BD, BK. Study Design and Data Discussion; BD, MMH, BK. Writing – review and editing; SG, VR, BD, MMH, BK. All authors approved the final version of the manuscript and have agreed to be personally accountable for their respective contributions and ensure that questions related to the accuracy or integrity of any part of the work, even those in which the author was not personally involved, are appropriately investigated, resolved, and the resolution documented in the literature.

## 8 Acknowledgements

We thank all staff at Biological Services Unit for their continuous effort in supporting our animal research with the highest standards of animal care and husbandry. In addition, the smooth operation of the TBCL3 facility and upholding safe working practices in the TB-high containment facility.

## References

Be, N. A., Lamichhane, G., Grosset, J., Tyagi, S., Cheng, Q. J., Kim, K. S., Bishai, W. R. & Jain, S. K. 2008. Murine model to study the invasion and survival of Mycobacterium tuberculosis in the central nervous system. J Infect Dis, 198, 1520–8.

Burn, C. G. & Finley, K. H. 1932. The Role of Hypersensitivity in the Production of Experimental Meningitis : I. Experimental Meningitis in Tuberculous Animals. J Exp Med, 56, 203–21.

Chan, E. D., Verma, D. & Ordway, D. J. 2020. Animal Models of Mycobacteria Infection. Curr Protoc Immunol, 129, e98.

Chin, J. H. 2014. Tuberculous meningitis: Diagnostic and therapeutic challenges. Neurol Clin Pract, 4, 199–205.

Davis, A. G., Donovan, J., Bremer, M., Van Toorn, R., Schoeman, J., Dadabhoy, A., Lai, R. P. J., Cresswell, F. V., Boulware, D. R., Wilkinson, R. J., Thuong, N. T. T., Thwaites, G. E., Bahr, N. C. & TUBERCULOUS MENINGITIS INTERNATIONAL RESEARCH, C. 2020. Host Directed Therapies for Tuberculous Meningitis. Wellcome Open Res, 5, 292.

Davis, A. G., Rohlwink, U. K., Proust, A., Figaji, A. A. & Wilkinson, R. J. 2019. The pathogenesis of tuberculous meningitis. J Leukoc Biol, 105, 267–280.

Du Preez, K., Seddon, J. A., Schaaf, H. S., Hesseling, A. C., Starke, J. R., Osman, M., Lombard, C. J. & Solomons, R. 2019. Global shortages of BCG vaccine and tuberculous meningitis in children. Lancet Glob Health, 7, e28–e29.

Hernandez Pando, R., Aguilar, D., Cohen, I., Guerrero, M., Ribon, W., Acosta, P., Orozco, H., Marquina, B., Salinas, C., Rembao, D. & Espitia, C. 2010. Specific bacterial genotypes of Mycobacterium tuberculosis cause extensive dissemination and brain infection in an experimental model. Tuberculosis (Edinb), 90, 268–77.

HOME OFFICE, U. 26 Oct 2022. Animals in Science Regulation Unit. https://www.gov.uk/government/collections/animals-in-science-regulation-unit.

Huynh, J., Donovan, J., Phu, N. H., Nghia, H. D. T., Thuong, N. T. T. & Thwaites, G. E. 2022. Tuberculous meningitis: progress and remaining questions. Lancet Neurol, 21, 450–464.

Kasahara, M. 1924. The production of tuberculosis meningitis in the rabbit and the changes in its cerebrospinal fluid. The American Journal of Diseases of Children, 27 (5), 428 –432.

Khatri, B., Keeble, J., Dagg, B., Kaveh, D. A., Hogarth, P. J. & Ho, M. M. 2021. Efficacy and immunogenicity of different BCG doses in BALB/c and CB6F1 mice when challenged with H37Rv or Beijing HN878. Sci Rep, 11, 23308.

Lee, J., Ling, C., Kosmalski, M. M., Hulseberg, P., Schreiber, H. A., Sandor, M. & Fabry, Z. 2009. Intracerebral Mycobacterium bovis bacilli Calmette-Guerin infection-induced immune responses in the CNS. J Neuroimmunol, 213, 112–22.

Mangtani, P., Abubakar, I., Ariti, C., Beynon, R., Pimpin, L., Fine, P. E., Rodrigues, L. C., Smith, P. G., Lipman, M., Whiting, P. F. & Sterne, J. A. 2014. Protection by BCG vaccine against tuberculosis: a systematic review of randomized controlled trials. Clin Infect Dis, 58, 470–80.

Manwaring, W. H. 1912. The Effects of Subdural Injections of Leucocytes on the Development and Course of Experimental Tuberculous Meningitis. J Exp Med, 15, 1–13.

Marais, S., Pepper, D. J., Schutz, C., Wilkinson, R. J. & Meintjes, G. 2011. Presentation and outcome of tuberculous meningitis in a high HIV prevalence setting. PLoS One, 6, e20077.

Ordway, D. J. & Orme, I. M. 2011. Animal models of mycobacteria infection. Curr Protoc Immunol, Chapter 19, Unit19 5.

Rich, A. R., Mccordock, H.A. 1933. The pathogenesis of tubercular meningitis. Bull John Hopkins Hosp, 52, 5–13.

Seddon, J. A., Wilkinson, R., Van Crevel, R., Figaji, A., Thwaites, G. E. & TUBERCULOUS MENINGITIS INTERNATIONAL RESEARCH, C. 2019. Knowledge gaps and research priorities in tuberculous meningitis. Wellcome Open Res, 4, 188.

Shope, R. E. & Lewis, P. A. 1929. A Paralytic Disease of Guinea Pigs Due to the Tubercle Bacillus. J Exp Med, 50, 365–70.

Skerry, C., Pokkali, S., Pinn, M., Be, N. A., Harper, J., Karakousis, P. C. & Jain, S. K. 2013. Vaccination with recombinant Mycobacterium tuberculosis PknD attenuates bacterial dissemination to the brain in guinea pigs. PLoS One, 8, e66310.

Tsenova, L., Ellison, E., Harbacheuski, R., Moreira, A. L., Kurepina, N., Reed, M. B., Mathema, B., Barry, C. E., 3RD & Kaplan, G. 2005. Virulence of selected Mycobacterium tuberculosis clinical isolates in the rabbit model of meningitis is dependent on phenolic glycolipid produced by the bacilli. J Infect Dis, 192, 98–106.

Tsenova, L., Harbacheuski, R., Moreira, A. L., Ellison, E., Dalemans, W., Alderson, M. R., Mathema, B., Reed, S. G., Skeiky, Y. A. & Kaplan, G. 2006. Evaluation of the Mtb72F polyprotein vaccine in a rabbit model of tuberculous meningitis. Infect Immun, 74, 2392–401.

WHO 27 Oct 2022. Global tuberculosis report 2022. https://www.who.int/publications/i/item/9789240061729.

Wilkinson, R. J., Rohlwink, U., Misra, U. K., Van Crevel, R., Mai, N. T. H., Dooley, K. E., Caws, M., Figaji, A., Savic, R., Solomons, R., Thwaites, G. E. & TUBERCULOUS MENINGITIS INTERNATIONAL RESEARCH, C. 2017. Tuberculous meningitis. Nat Rev Neurol, 13, 581–598.

Zucchi, F. C., Tsanaclis, A. M., Moura-Dias, Q., JR., Silva, C. L., Pelegrini-Da-Silva, A., Neder, L. & Takayanagui, O. M. 2013. Modulation of angiogenic factor VEGF by DNA-hsp65 vaccination in a murine CNS tuberculosis model. Tuberculosis (Edinb), 93, 373–80.

